# T1 white/gray contrast as a predictor of chronological age, and an index of cognitive performance

**DOI:** 10.1101/171892

**Authors:** John D. Lewis, Alan C. Evans, Jussi Tohka, for the Brain Development Cooperative Group, and the Pediatric Imaging, Neurocognition, and Genetics Study

## Abstract

The maturational schedule of human brain development appears to be narrowly confined. The chronological age of an individual can be predicted from brain images with considerable accuracy, and deviation from the typical pattern of brain maturation has been related to cognitive performance. Methods using multi-modal data, or complex measures derived from voxels throughout the brain have shown the greatest accuracy, but are difficult to interpret in terms of the biology. Measures based on the cortical surface(s) have yielded less accurate predictions, suggesting that perhaps developmental changes related to cortical gray matter are not strongly related to chronological age, and that perhaps development is more strongly related to changes in subcortical regions or in deep white matter. We show that a simple metric based on the white/gray contrast at the inner border of the cortical gray-matter is a comparably good predictor of chronological age, and our usage of an elastic net penalized linear regression model reveals the brain regions which contribute most to age-prediction. We demonstrate this in two large datasets: the NIH Pediatric Data, with 832 scans of typically developing children, adolescents, and young adults; and the Pediatric Imaging, Neurocognition, and Genetics data, with 760 scans of individuals in a similar age-range. Moreover, we show that the residuals of age-prediction based on this white/gray contrast metric are more strongly related to IQ than are those from cortical thickness, suggesting that this metric is more sensitive to aspects of brain development that reflect cognitive performance.

## 1. Introduction

Human brain development begins in the third gestational week and extends at least into adulthood, and arguably to the end of life (Stiles and Jernigan, 2010). A neonate has approximately the adult cortical layer structure and cortical folding pattern, and the major fiber pathways are complete; but, the brain almost quadruples in size during early childhood (Dekaban and Sadowsky, 1978; Pfefferbaum et al., 1994), and there are vast changes in the composition of both gray and white matter. Such changes continue throughout development and senescence, albeit at a slower pace. Trajectories of maturation are region-dependent. Brain regions associated with sensory and motor processes mature early; secondary processing regions mature over childhood and adolesence; association areas continue to develop into adulthood (Casey et al., 2005). The timing of these developmental brain changes appears to be precisely controlled. Chronological age can be predicted from brain images with reasonable accuracy, even setting aside the dramatic changes of prenatal and early postnatal development. Correlations between chronological age and age estimates from brain images are as high as 0.93 based on a single modality (Franke et al., 2012); and as high as 0.96 based on multi-modal data (Brown et al., 2012).

Moreover, differences in brain maturational trajectories are associated with differences in cognition. Developmental disorders are generally associated with accelerated or delayed patterns of brain maturation (Toga et al., 2006). And, within the bounds of typical development, differences in the trajectories of cortical gray-matter maturation throughout childhood and adolesence are associated with differences in intelligence scores (Shaw et al., 2006), and in the effectiveness of executive functions (Gogtay et al., 2004). In terms of MRI-based age predictions, the residuals of predictions have been associated with cognitive processing speed (Erus et al., 2015), and with discrepancies in verbal and performance IQ (Khundrakpam et al., 2015). Thus, divergence of brain-age predictions from chronological age appears to index cognitive precocity or delay (Franke et al., 2012; Bunge and Whitaker, 2012; Erus et al., 2015).

In addition to knowing whether an individual is showing precocious or delayed development, there is value in knowing what generates that assessment. Previous literature leaves that quite unclear. Brain maturation appears to be reflected in a broad array of changes in all modalities considered. Estimates of age from voxel-based morphometry yield a correlation with chronological age of 0.93, but it is not clear what microstructural changes underlie these estimates (Franke et al., 2012). Estimates of age from T2-weighted signal intensity show a correlation with chronological age of 0.91, with contributions from both white-matter tracts and subcortical gray (Brown et al., 2012). Estimates of age involving diffusion data show a correlation close to 0.9, with the greatest contributions coming from long-range connections, e.g. the corpus callosum, and from both cortical and subcortical gray matter (Mwangi et al., 2013; Erus et al., 2015; Brown et al., 2012). Estimates of age based on cortical thickness show a correlation of 0.84, with the greatest contribution to predictions coming from sensori-motor and association areas (Khundrakpam et al., 2015). Estimates of age based on resting state functional connectivity show a correlation of 0.75, with the greatest contribution to predictions associated with segregation into sub-networks (Dosenbach et al., 2010). And the multi-modal results of Brown et al. (2012) yield a correlation of 0.96, with the greatest relative contributions to the predictions coming from subcortical T2 signal intensity, diffusion measures in white-matter tracts, and from subcortical volumetric and diffusion measures. Taken together, these results suggest a surprisingly limited role for cortical measures in age prediction.

The age-predictions based on cortical thickness (Khundrakpam et al., 2015) were far inferior to those based on white-matter diffusion measures or T2 signal intensity (Mwangi et al., 2013; Erus et al., 2015; Brown et al., 2012), or to results from measures that included subcortical material (Brown et al., 2012); and the the multi-modal analysis of Brown et al. (2012) showed the relative contributions of both cortical thickness and cortical surface area to be low. And yet, the best single modality predictions were derived from T1-weighted data (Franke et al., 2012), albeit from whole-brain measures.

Conceivably, the signal that drives the exceptional prediction performance of Franke et al. (2012) comes neither from the subcortical material nor from the white-matter nor the cortex, but from the differences between the white matter and the gray matter. One of the most salient features of T1-weighted images is the intensity contrast between gray- and white-matter. This contrast has been measured as the ratio of the signal intensity on either side of the surface formed at the gray-white cortical boundary, and has been shown to be more strongly correlated with age in many cortical regions than is cortical thickness (Salat et al., 2009). Indeed, in neonates, intensity contrast is reversed: the cortico-cortical connections appear darker than the cortical gray-matter, whereas in adults, as a result of myelination, these cortico-cortical connections appear white. Moreover, this intensity contrast has proven useful in characterizing typical patterns of brain development (Salat et al., 2009; Westlye et al., 2010), and identifying brain abnormalities in developmental disorders such as autism spectrum disorder (Andrews et al., 2017) as well as disorders primarily afflicting the elderly such as Alzheimer’s disease (Salat et al., 2011; Jefferson et al., 2015). Thus, we tested the hypothesis that intensity contrast at the gray-white cortical boundary will allow for more accurate age predictions than cortical thickness, and perhaps better insight into the relation between brain maturation and cognitive development.

## 2. Materials and Methods

### 2.1. Data

The data used were taken from two large-scale datasets: the NIH Pediatric data (NIHPD); and the Pediatric Imaging, Neurocognition, and Genetics data (PING). Detailed descriptions of the NIHPD and PING samples are given in (Evans et al., 2006) and (Brown et al., 2012), respectively; here we give a brief summary. Table 1 provides the demographic information for the two populations.

**Table 1:** Subject demographics. Ages are given in years.

|  | Subjs (Males) | Scans (Males) | Mean age | Min age | Max age |
| --- | --- | --- | --- | --- | --- |
| NIHPD | 401 (191) | 832 (389) | 12.5 | 4.88 | 22 |
| PING | 760 (398) | 760 (398) | 12.3 | 3.17 | 21 |

#### 2.1.1. The NIHPD sample

The NIHPD data were collected to allow the characterization of healthy brain development, and the relation between brain development and behaviour.

Both high-quality MRI data and comprehensive clinical/behavioral measures were collected. Recruitment was epidemiologically based, demographically balanced, and used strict exclusion factors. The sample includes over 400 children ranging from 4.5 to 18.5 years of age, well-distributed across the range. The database is a mix of longitudinal and cross-sectional data, but we consider all data as cross-sectional, taking care to ensure that the evaluation of predictive models is not biased by this dependence. The data were collected at six sites: Children’s Hospital, Boston; Children’s Hospital Medical Center, Cincinnati; University of Texas Houston Medical School, Houston; Neuropsychiatric Institute and Hospital, UCLA; Children’s Hospital of Philadelphia; and Washington University, St. Louis. Data were acquired on either a General Electric 1.5T scanner or a Siemens Medical Systems 1.5T scanner. Each site acquired multiple contrast data (T1-weighted, T2-weighted and PD-weighted); the current study utilizes only the T1-weighted data.

#### 2.1.2. The PING sample

The PING dataset includes data from more than 800 typically developing children between the ages of 3 and 20 years, including individuals with learning or language disabilities. Subjects were excluded if they had a history of major developmental, psychiatric, and/or neurological disorders, or medical conditions that affect neurological development, or had had a brain injury. As opposed to the NIHPD sample, these data are strictly cross-sectional.

Data were collected at 10 sites: Weil Cornell Medical College, University of California at Davis, University of Hawaii, Kennedy Krieger Institute, Massachusetts General Hospital, University of California at Los Angeles, University of California at San Diego, University of Massachusetts Medical School, University of Southern California, and Yale University. Data were acquired on either a General Electric 3T scanner, a Siemens Medical Systems 3T scanner, or a Philips 3T scanner. Across sites and scanners, a standardized multiple modality high-resolution structural MRI protocol was implemented. Each site acquired T1-, T2-, and diffusion-weighted scans; the current study uses only the T1-weighted data.

### 2.2. Surface measurements

#### 2.2.1. Surface extraction

The T1-weighted volumes were processed with CIVET (version 2.1), a fully automated structural image analysis pipeline developed at the Montreal Neurological Institute. CIVET corrects intensity non-uniformities using N3 (Sled et al., 1998); aligns the input volumes to the Talairach-like ICBM-152-nl template (Collins et al., 1994); classifies the image into white matter, gray matter, cerebrospinal fluid, and background (Zijdenbos et al., 2002; Tohka et al., 2004); extracts the white-matter and pial surfaces (Kim et al., 2005); and maps these to a common surface template (Lyttelton et al., 2007).

#### 2.2.2. Cortical thickness measurements

Cortical thickness (CT) is measured in native space at 81,924 vertices using the Laplacian distance between the two surfaces. The Laplacian distance is the length of the path between the gray and white surfaces following the tangent vectors of the cortex represented as a Laplacian field (Jones et al., 2000). This ensures that any misalignment of the vertices in the white and gray surfaces do not introduce error in the thickness measures.

#### 2.2.3. White/gray contrast measurements

To extract the white/gray contrast measures, the intensity on the T1-weighted MRI was sampled 1 mm inside and 1 mm outside the white surface, and the ratio of the two measures was formed. More precisely, a distance map was created from the white surface at 0.25mm resolution; the distance map was smoothed with a 0.5 mm FWHM Gaussian kernel; and a gradient vector field was computed. The white surface was then moved 1mm inward along this gradient vector field to produce a sub-white surface, and 1mm outward to produce a supra-white surface. The T1-weighted intensity values were sampled at each vertex of both the supra-white surface and the sub-white surface. Then the ratio was formed by dividing the value at each vertex of the sub-white surface by the value at the corresponding vertex of the supra-white surface. This procedure is depicted in Figure 1.

**Figure 1:**
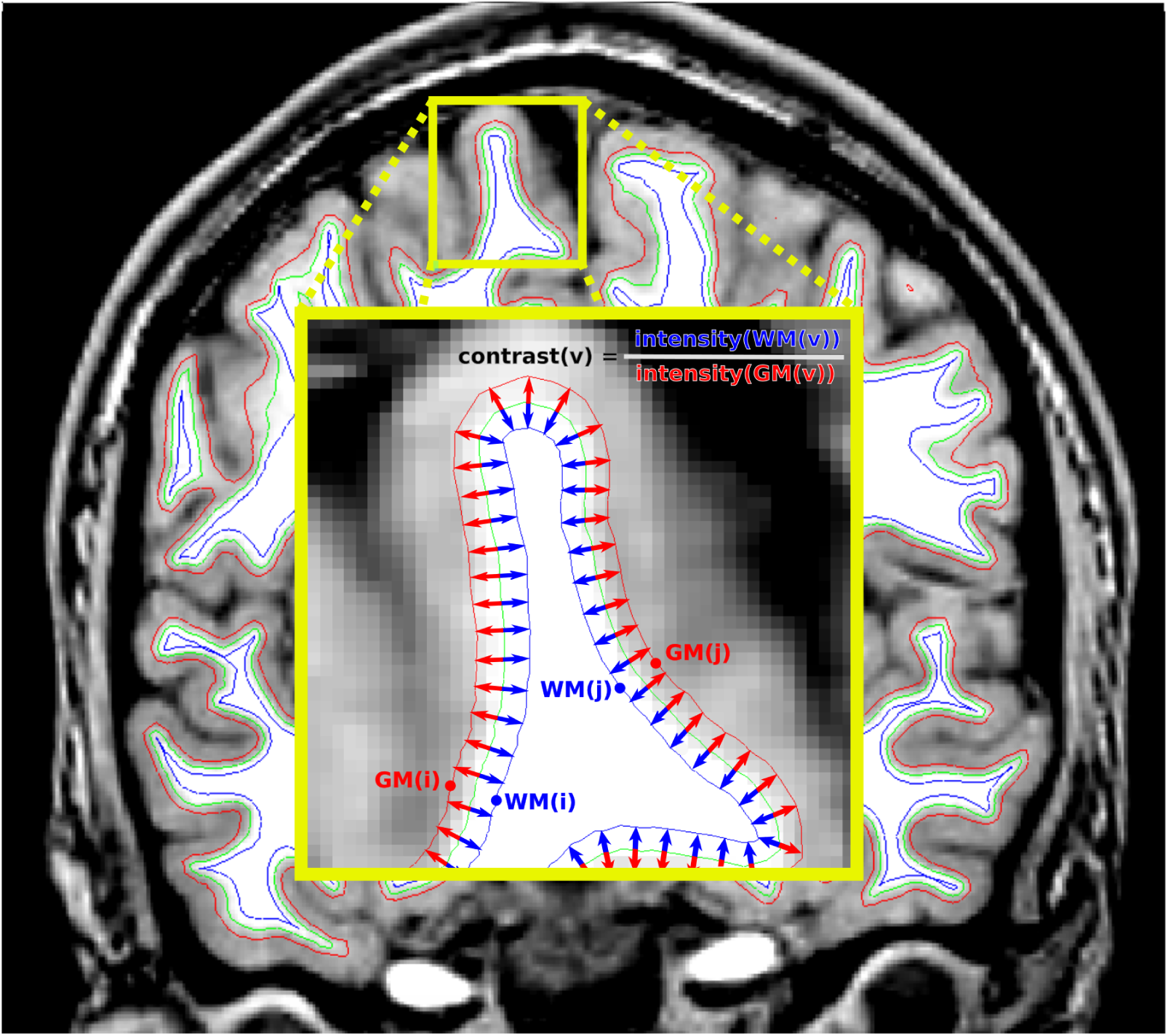
The surface at the gray-white border (green) is extracted by CIVET. A distance map and the corresponding vector field were created from this, and copies of the surface were moved outward 1 mm (red) and inward 1 mm (blue) along the gradient vectors of this map. The magnified inset illustrates this procedure. Gray matter and white matter intensity were measured at the vertices of the surface 1mm out and 1 mm in, respectively. The white/gray contrast measure is the ratio of the white intensity to gray intensity.

### 2.3. Age prediction

Our age prediction method is adapted from (Khundrakpam et al., 2015). We assumed a linear model for predicting subjects’ ages based on cortical measurements. The model is

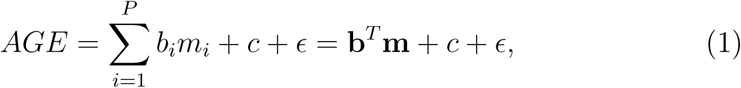

where AGE is the age of the subject (in days); *m_i_*, *i* = 1 …*P*, are the cortical measurements (cortical thickness or white/gray contrast); *b_i_* and *c* are the model weights to be learned by the machine learning algorithm, and *ϵ* is an error term. We also considered age prediction with cortical thickness and white/gray contrast where the two types of cortical measurements are concatenated yielding a model with two measurements at each location. We standardized the variables *m_i_* so that each of them has unit variance and zero mean as this is important for the interpretation of the prediction model. We considered several spatial resolutions of the measurements. The original 81,924 measurements on the cortical surface were grouped into smaller sets and averaged. The number of parcels chosen were 78, 160, 640, and 2560, thus *P* = 78,160, 640, 2560 for a single measurement type and *P* = 156, 320,1280, 5120 for the combined measurements. The 78 parcel case was obtained by averaging cortical measurements in each cortical region of the Automated Anatomical Labelling (AAL) atlas (Tzourio-Mazoyer et al., 2002), a commonly used atlas in analyses of cortical measures. The other cases were not based on an atlas, but rather were obtained by recursively merging the neighbouring triangles of the surface mesh model.

We estimated the parameters b = [*b*_1_,…, *b_P_*] and *c* of the model by penalized least squares with the elastic-net penalty as implemented in the Glmnet package (Friedman et al., 2010). This equals to the minimization of the cost function

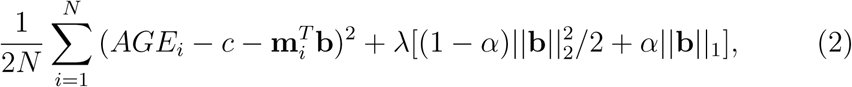

where subscript *i* refers to scans, **m**_*i*_ are the measurements for subject *i* and *N* is the number of scans. The elastic-net penalty (1 – *α*)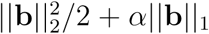 is a weighted average of the LASSO and ridge penalties (Zou and Hastie, 2005). The LASSO penalty ||**b**||_1_ forces many parameters to have a zero-value leading to the variable selection while the ridge penalty 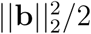 helps to ensure that highly correlated variables are selected simultaneously. We selected the two penalties to have an equal weight, i.e. set *α* = 0.5, as in (Khundrakpam et al., 2015). The relative weight of the data term and the penalties, denoted by the parameter λ ∈ ℝ in Eq. (2), was decided based on cross-validation from a sequence of 300 values decreasing on the log scale (Friedman et al., 2010). We used this algorithm as a baseline method due to its good performance in (Khundrakpam et al., 2015) and attractive properties for brain imaging (Carroll et al., 2009; Khundrakpam et al., 2015).

With the PING dataset, the white/gray contrast measures varied between the different scanner manufacturers. More specifically, the white/gray contrast measures from the Philips scanners were different from the measures by the Siemens and GE scanners. However, normalizing the contrast values per scanner manufacturer worked reasonably well as a correction. In more detail, the normalized contrast measure is 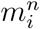 = (*m_i_* – *μ_scanner_*)*σ_scanner_*, where *μ_scanner_* and *σ_scanner_* are the mean and the standard deviation of the contrast values across whole brain and across all the subjects (see supplementary Figure S1 for details). We did not encounter the problem with the NIHPD dataset, which was acquired with Siemens and GE scanners. However, to maintain methodological consistency the same normalization was applied also to the NIHPD data. With cortical thickness measures, the normalization decreased the predictive accuracy and was not applied.

### 2.4. Evaluation of age prediction

We selected the λ and evaluated the age predictions using two nested stratified 10-fold cross-validation (CV) loops (Ambroise and McLachlan, 2002; Huttunen et al., 2012) ^1^. The stratification of the CV means here that the age distribution of the subjects in each of the 10 CV folds was approximately the same. The value for the parameter λ was selected in the inner CV loop by minimizing the mean squared error among the candidate values, and the age predictions were evaluated in the outer CV loop, thus avoiding the training on testing data problem. We repeated the nested CV loops 10 times to reduce the random variation in the evaluation of accuracy due to the random selection of the folding scheme. The model goodness criteria applied were the correlation coefficient between the chronological and estimated age, and the mean absolute error (MAE) between the chronological and estimated age. For the correlation coefficient, we averaged the 10 distinct correlation values stemming from the 10 CV runs. Since with the NIHPD sample we had up to three cortical measurements of certain subjects at different ages, we controlled for the non-independence of the observations in the CV by placing all the scans of the subject *i* in the same fold and thus all the scans were either in the training set or in the test set. Therefore, data from subject *i* was never used to build an age-predictor for subject *i*. This is an important consideration and failure to account for non-independence would lead to positively biased model accuracy estimates.

We computed 95% confidence intervals (CIs) for cross-validated correlations using a bootstrap method similar to the standard bootstrap applied to the correlation coefficients and detailed in the appendix. These CIs approximate the generalization performance of the prediction measured by cross-validated correlations. Likewise, we compared the cross-validated MAEs from two different measurements using a permutation test adapting the same resampling scheme as the bootstrap method detailed in the appendix ^2^.

With the NIHPD database there were IQ scores associated with a subsample of 760 scans from 391 subjects; we studied whether the residuals from the age predictions would be informative in explaining the IQ of the children. We define the age residual as chronological age minus predicted age. Again, we considered the data as cross-sectional, taking care to ensure that the longitudinal data did not bias the study. We constructed linear regression models explaining different IQ scores (full scale IQ (FSIQ), verbal IQ (VIQ) and performance IQ (PIQ)) in terms of the age residuals, controlling for chronological age, gender, and total brain volume, which are known to correlate with aspects of IQ. This experiment was repeated for all three types of age residuals (based on thickness, contrast, and thickness+contrast) and for all parcellations. For each type of age residual, we selected the predicted age of each subject to be the average across 10 cross-validation runs. We recorded the t-score of the regression coefficient corresponding to the age residual in the model and the adjusted R-squared of the model. To test the hypothesis that the model with the age residual based on measurement A (e.g., white/gray contrast) is better at explaining the IQ than the model with the age residual based on measurement B (e.g., thickness), we used the J-test for non-nested models (Watnik et al., 2001; Davidson and MacKinnon, 1981). The comparison of two non-nested models is challenging (Greene, 2003) and the J-test is a two-step procedure testing two hypotheses (model A is better than model B, and vice versa). All four possible outcomes can occur (reject both hypotheses, reject neither, or either one) and the test is inconclusive if both hypotheses are rejected or neither of them are rejected (Greene, 2003).

## 3. Results

### 3.1. Age prediction accuracy

The cross-validated accuracies of the age predictions are reported in Table 2 and plotted in Figure 2 with significant differences in mean absolute errors indicated. The white/gray contrast measures yielded significantly more accurate predictions than the cortical thickness measures with every parcellation and with both databases. The white/gray contrast and cortical thickness measures together resulted in more accurate age predictions than either type of measure alone.

**Table 2:** The cross-validated accuracies of age predictions based on cortical thickness, white/gray contrast, or both measures together. The average cross-validated correlation value (*R*) for each is given, as well as the mean absolute error (*MAE*) in days.

|  |  | Thickness |  | Contrast |  | Contrast+Thickness |  |
| --- | --- | --- | --- | --- | --- | --- | --- |
| | parcels | $R$ | $MAE$ | $R$ | $MAE$ | $R$ | $MAE$ |
| NIHPD | AAL | 0.77 | 728 | 0.83 | 633 | 0.88 | 535 |
|  | 160 | 0.81 | 681 | 0.86 | 575 | 0.89 | 514 |
|  | 640 | 0.82 | 652 | 0.88 | 547 | 0.89 | 504 |
|  | 2560 | 0.81 | 671 | 0.87 | 551 | 0.89 | 510 |
| PING | AAL | 0.86 | 738 | 0.89 | 684 | 0.92 | 560 |
|  | 160 | 0.87 | 724 | 0.90 | 661 | 0.92 | 561 |
|  | 640 | 0.88 | 675 | 0.91 | 619 | 0.92 | 553 |
|  | 2560 | 0.89 | 656 | 0.91 | 600 | 0.92 | 558 |

**Figure 2:**
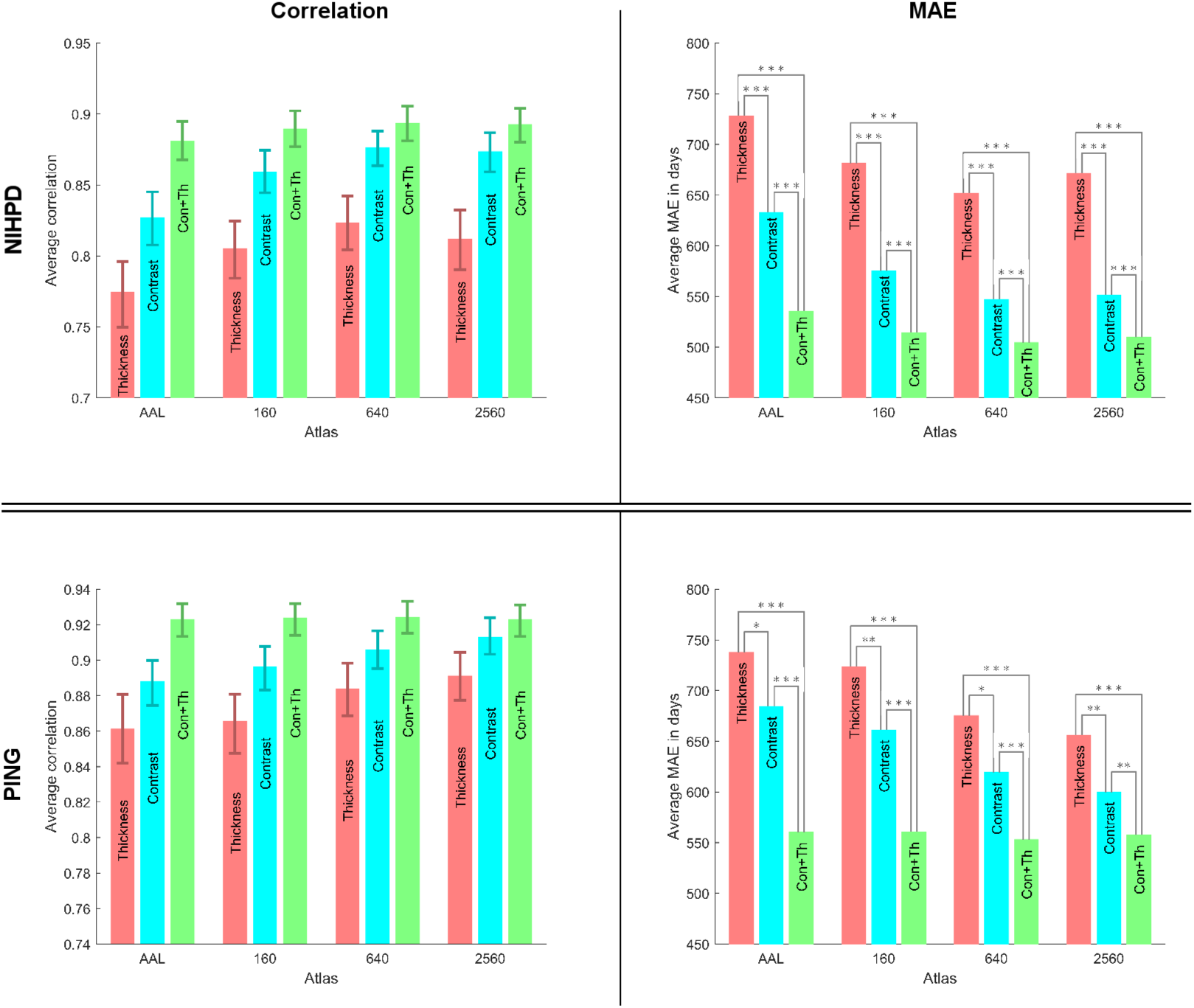
The accuracies of the age predictions using different cortical measures. Glmnet was used as the machine learning algorithm. The correlation plots show, in addition to the average cross-validated correlation value, the 95 % confidence intervals obtained by a bootstrap method. The MAE plots show, in addition to the average cross-validated MAE, a permutation test based comparison between the different methods: n.s stands for not significant, ^*^ for *p* < 0.05 ^**^ for *p* < 0.01 and ^***^ for *p* < 0.001.

Figure 3 shows scatter-plots of the predicted versus chronological age during a single cross-validation run (one out of ten). It is interesting to note that the age estimates varied considerably between the different cross-validation runs. The average absolute deviations (AADs) varied from 58 days (with the white/gray contrast measure and the AAL atlas) to 134 days (with cortical thickness and the 2560 parcel atlas). Generally, the AAD increased with the atlas resolution, and the lowest AADs at a given resolution were with the white/gray contrast measure. The plots in Figure 3 are qualitatively similar, but thickness shows greater variance than do white/gray contrast or the combination of measures. Gender and scanner manufacturer had no observable effect on the direction or the magnitude of the residuals.

**Figure 3:**
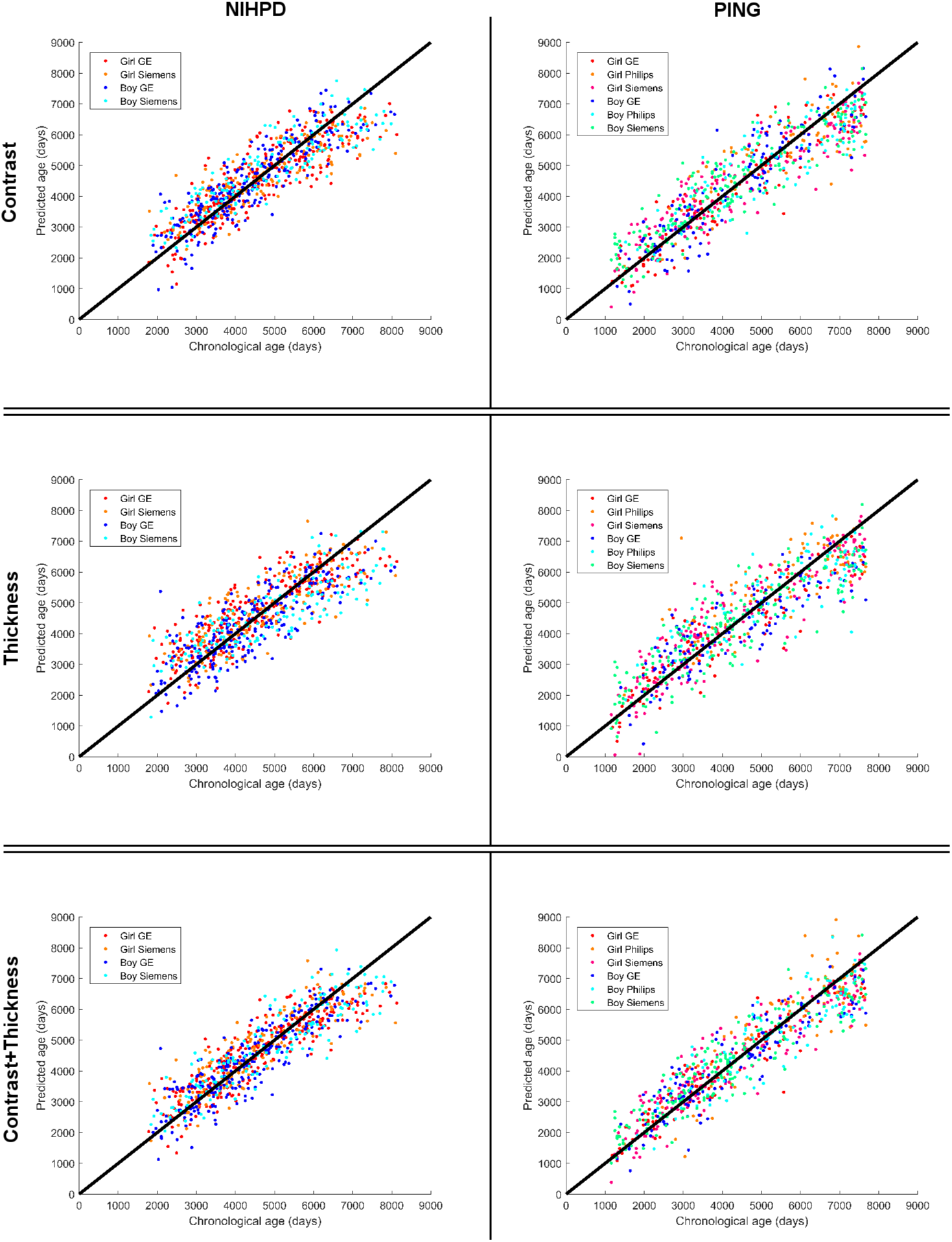
Scatter-plots of the predicted versus chronological age during a single crossvalidation run (one out of ten) with 640 parcels. The black line has a slope of 1 and originates at the origin; this line depicts the optimal predictions. The scanner manufacturer/gender combinations are shown with different colors. No apparent bias either due to gender or scanner manufacturer can be observed. Note the lesser spread of the predictions with white/gray contrast than with cortical thickness, and the lesser spread of the predictions with the combined measures than with either alone.

### 3.2. Age predictors

Our usage of an elastic net penalized linear regression model (Friedman et al., 2010) reveals the brain regions which contribute most to age-prediction. The cortical parcels for which the white/gray contrast or thickness measures reliably contribute to age prediction are shown in Figure 4, for both databases. We defined the signed importance as the median value of weight of the parcel *i* (*b_i_*) in the linear regression model across the 10 x 10 cross-validation; thus a parcel had to be selected in the age model at least 50 times during the 10 x 10 cross-validation runs to achieve a non-zero value. The figure shows the non-zero signed importances of all parcels for both measures for both databases. The figure uses the 640 parcel model, as this resolution yielded the highest accuracy in the majority of cases (see Figure 2). Both white/gray contrast and thickness showed similar predictor importance patterns across both databases. Both measures showed predictor importances broadly distributed across cortex, meaning that the age predictions were constructed from information throughout the cortex.

**Figure 4:**
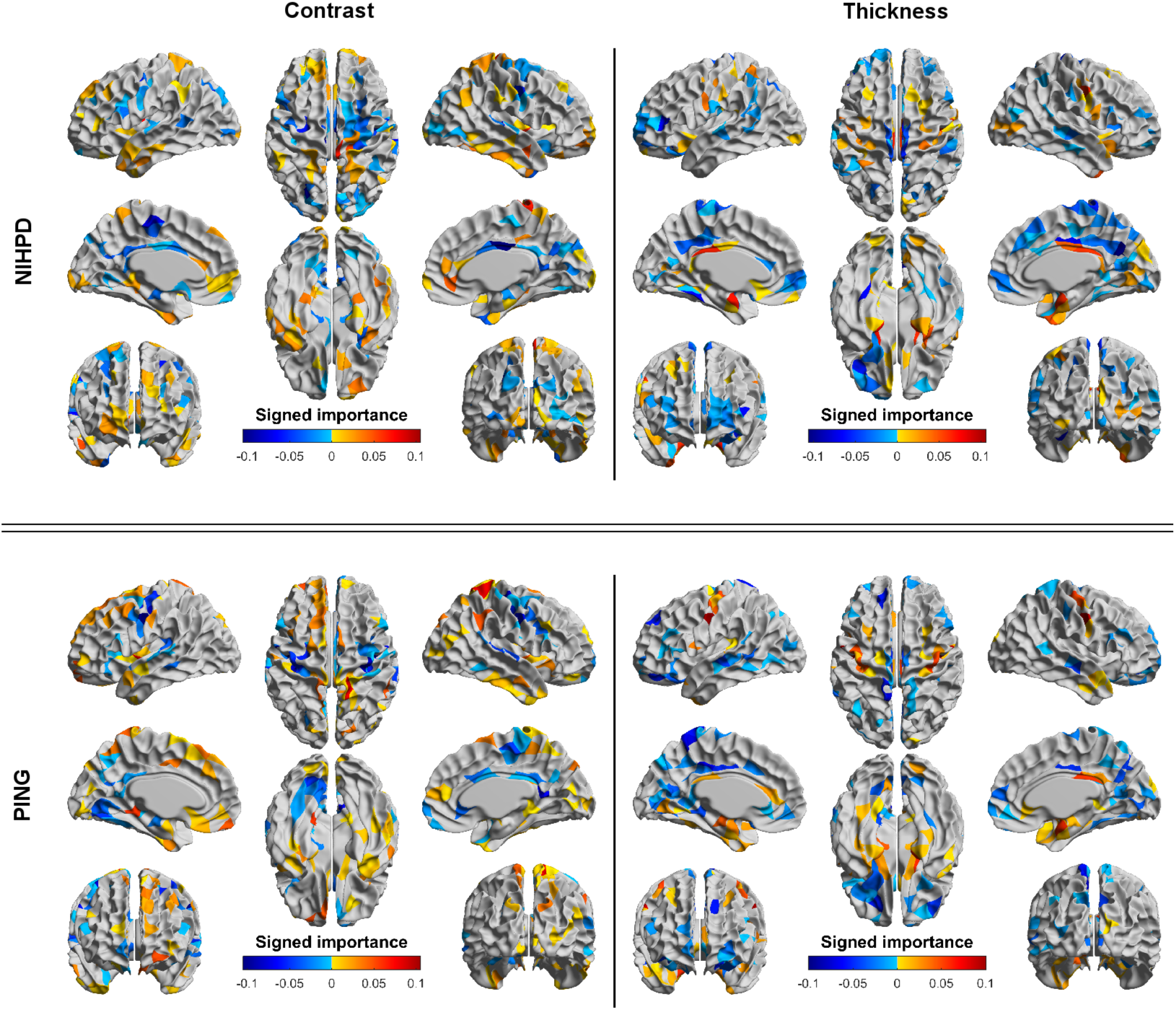
The signed importance of different parcels for age predictions for white/gray contrast and for thickness in both datasets with the 640 parcel model. The signed importance is defined as the median value of weight *b_i_* across the 10 x 10 cross-validation; thus a parcel has to be selected in the age model at least 50 times during the 10 x 10 cross-validation runs to show a non-zero value in the plots. The weights were computed using standardized data so their values are comparable across the cortex. The top row shows the signed importances in the NIHPD data; the bottom row shows the signed importances in the PING data. The left column shows the signed importances with white/gray contrast; the right column shows the signed importances with thickness.

But, for both white/gray contrast and for thickness, the spatial distributions of the positive versus negative signed importances indicate that different parts of cortex contribute different information. For white/gray contrast, regions involved in low-level processing, e.g. auditory, somatosensory, and motor cortex, tended to show negative signed importances, and association cortices tended to show positive signed importances. For thickness, this pattern was reversed.

### 3.3. Age prediction residuals and IQ

We analyzed the relation between cognitive performance and errors in age prediction. We defined the age residual as chronological age minus predicted age, and looked at its relation to IQ. There were IQ scores associated with a subsample of 760 scans from 391 subjects from the NIHPD dataset; PING does not have IQ measures, and so was not included in the analysis. We constructed linear regression models explaining different IQ scores (full scale IQ (FSIQ), verbal IQ (VIQ) and performance IQ (PIQ)) in terms of the age residuals, controlling for chronological age, gender, and total brain volume, which are known to correlate with aspects of IQ. The key statistics of the regression models explaining IQ are shown in Table 3. The models with the age residuals based on the contrast measures yielded the highest *R*^2^ in all the cases. We used the J-test for non-nested models (Watnik et al., 2001; Davidson and MacKinnon, 1981) to statistically assess the age models based on different cortical measures. According to the J-tests, the models with the age residuals based on the contrast measures alone were more appropriate than others, except in two cases in which the tests were inconclusive between the contrast measures and the combined measures (for VIQ, with either the AAL parcels or the 2560 parcels). The age residuals based on the contrast measures alone were significant in all the regression models except the one modeling VIQ using the AAL parcels. Likewise, the age residuals based on the combined measures were significant for FSIQ and PIQ, and for VIQ with the highest resolution parcellation. These results differ starkly from those for the age residuals based on the thickness measures, which show no significant relationships with IQ.

**Table 3:**
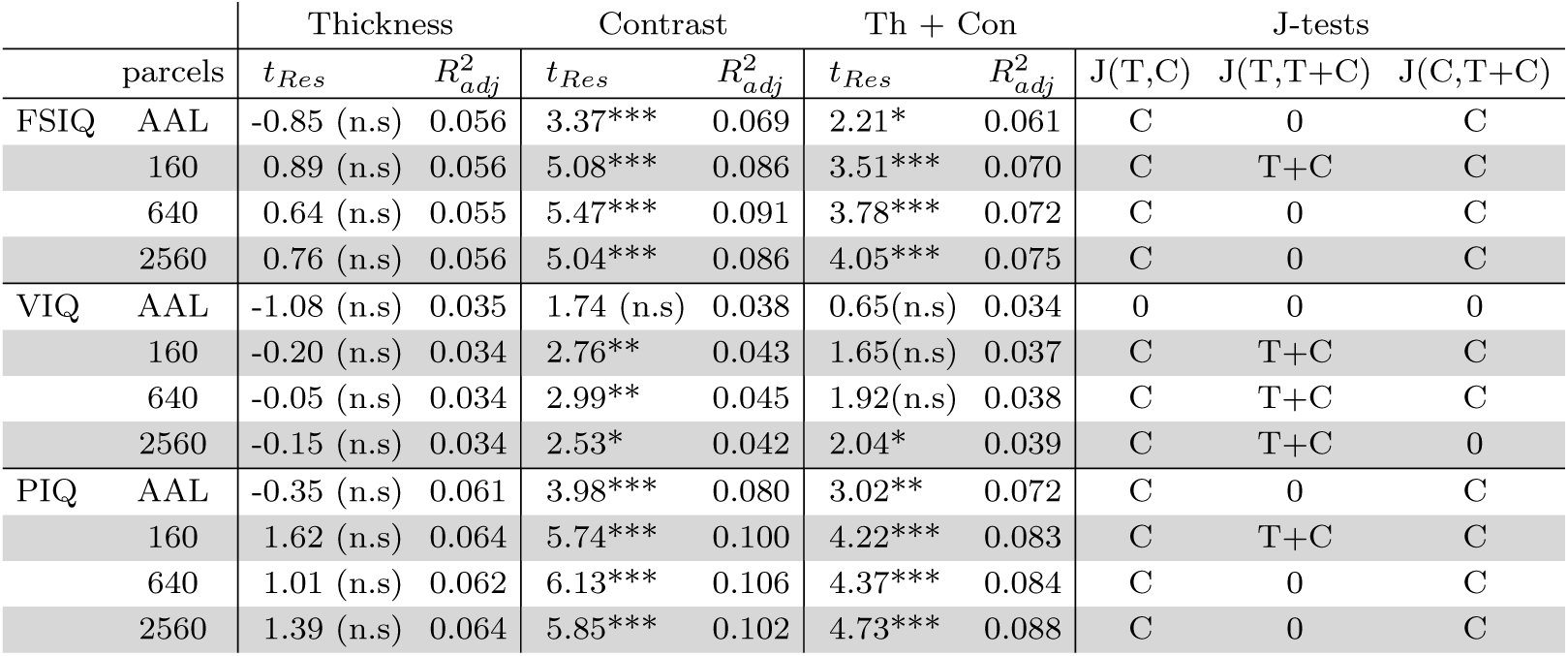
The key statistics of models explaining IQ. Age residual is defined as chronological age minus predicted age. *t_Res_* is the t-statistic corresponding to the age residual in the model. n.s stands for not significant, ^*^ for *p* < 0.05 ^**^ for *p* < 0.01 and ^***^ for *p* < 0.001. 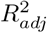 is the adjusted *R*^2^ of the whole model (the percentage of variance explained (adjusted)). J-Test columns indicate if either of the models was more appropriate in the pairwise comparison. *p* < 0.025 was used as the significance threshold. The columns list the more appropriate model (T for thickness, C for contrast, and T + C for combined thickness and contrast) or 0 if the test was inconclusive.

## 4. Discussion

Previous research has shown that chronological age can be predicted with high accuracy based on multiple measures from multi-modal data (Brown et al., 2012), or even from a single modality, but from complex measures derived from voxels throughout the brain (Franke et al., 2012). Measures based on the cortical surface(s), e.g cortical thickness or area, have yielded less accurate predictions (Brown et al., 2012; Khundrakpam et al., 2015), suggesting that perhaps developmental changes related to cortical gray matter are not strongly related to chronological age, and that perhaps development is more strongly related to changes in subcortical regions or in deep white matter. Here we have shown that a single measure based on the surface at the inner edge of the cortical gray matter, i.e. the white/gray contrast, yields comparable prediction accuracy, and prediction accuracy improves further with the inclusion of cortical thickness measures. Moreover, in partial accord with previous findings relating differences between chronological age and predicted age to cognitive functioning (Franke et al., 2012; Erus et al., 2015), we have shown that the residuals of age prediction based on this white/gray contrast are strongly related to IQ, particularly performance IQ.

Additionally, unlike methods that have used nonlinear mapping functions (i.e., kernels), our use of an elastic net penalized linear regression model allowed us to identify the cortical regions which contributed to our age predictions. Cortical regions in all lobes contributed to the predictions, both for measures of white/gray contrast and for measures of thickness, and for both measures together. But the signed importances of these contributions differed for the two measures. Measures of white/gray contrast tended to have negative signed importance in regions involved in low-level sensory processing, indicating a relative decrease with age, and positive signed importance in association cortex, indicating a relative increase with age; measures of thickness showed the opposite pattern, i.e. they tended to have positive signed importance in regions involved in low-level processing, and negative signed importance in association cortex.

The opposing nature of the signed importances for contrast and thickness is informative. It suggests that a portion of cortical thinning during development may not correspond to actual reductions in the thickness of the cortex, but rather may reflect increases in intracortical myelination and structure, or in the white matter adjacent to cortex, or both. Cortico-cortical connections typically originate in layer III and terminate in layers I and II. These fibers constitute the U-fibers that inter-connect adjacent gyri, as well as long-distance connections, including inter-hemispheric connections. Maturation of the U-fibers, which lie directly beneath the cortex, means increases in their diameter and myelination; maturation of cortico-cortical connectivity, in general, will increase intra-cortical myelination and cell complexity. These changes will impact both white/gray contrast and, to a lesser extent, measures of thickness. The effect on contrast is direct. The effect on thickness is due to the elongation of the gradient between white and gray matter, which determines the placement of the surface defining the inner edge of cortical gray matter; this effect will be more subtle. In addition, thalamocortical afferents ascend to cortical layer IV; and the efferents that project to the thalamus are in layer VI, with dendritic arbours that extend to layer IV. Maturation of these fibers will reduce white/gray contrast further and further elongate the gradient between the two. But, there is a greater prominence of thalamocortical afferents and efferents in primary sensory cortex than elsewhere. Thus the transition from white matter to gray matter in primary sensory cortex will become more clouded than elsewhere as these connections become more myelinated. This will be strongly reflected in reduced measures of contrast, which will show greater age-related reductions than the rest of cortex. Cortical thinning due to regressive processes will be opposed by the development of the dendritic arbours extending from layer VI to Layer IV, and further impacted by the more elongated gradient between white and gray matter produced by the combination of the invading cortico-cortical connection and subcortical-cortical connections. Hence the opposing nature of the signed importances for primary sensory regions versus association cortex, and the apparent developmental cortical thickening in primary sensory and motor regions, but cortical thinning in association areas (Lyall et al., 2015).

This pattern, of course, also reflects differences in developmental timing. Histological studies have shown that white-matter development is more rapid for fibers directly connected to primary sensory and motor regions than for fibers connecting secondary processing regions, and more prolonged for fibers connecting association areas (Yakovlev and Lecours, 1967). Additionally, diffusion tensor imaging over development, as well as longitudinal four-dimensional mapping of cortical and subcortical anatomy has shown that subcortical material develops over a more extended period of time than does cortex and cortico-cortical connectivity (Lebel et al., 2008; Raznahan et al., 2014). Thus, the early maturation of fibers connected to primary sensory areas will yield a more rapid decline in white/gray contrast in these regions than elsewhere, with the above-mentioned impact on cortical thickness measures. The more prolonged period of development of subcortical-cortical connections will further distinguish primary sensory and motor areas from other areas, particularly association areas.

Though the measures of white/gray contrast and cortical thickness are related in this opposing way, the two measures do not completely mirror each other. Age predictions based on contrast were considerably more accurate than those based on thickness, with an improvement in mean absolute error of approximately 16 percent in the NIHPD sample, and 8.5 percent in PING. But, it is also not the case that contrast is simply more accurate. Age predictions based on both contrast and thickness were more accurate than those based on contrast alone, showing an improvement of approximately 8 percent in the two samples. Thus there are age-related alterations in thickness that are not captured by white/gray contrast.

Seemingly paradoxically, the residuals of age prediction based on white/gray contrast relate more strongly to measures of cognition, i.e. FSIQ, VIQ, and PIQ, than do the residuals based on both contrast and thickness. But, plausibly this is due to the more limited variance in the residuals for the combined measures. The residuals of the less accurate predictions based on only cortical thickness, however, were unrelated to these cognitive measures (c.f. Khundrakpam et al., 2015). This was the case for each of the cognitive measures.

Interestingly, the residuals, defined as *chronological age* - *predicted age*, were positively correlated with the cognitive measures. Thus, the younger a brain appears relative to its actual age, the greater the intelligence. This appears to conflict with Erus et al. (2015), albeit their results are in a slightly older sample, with alternate cognitive measures. Erus et al. (2015) reported that individuals with brains that appeared younger than their their chronological age showed inferior cognitive processing speed, and vice-versa. This was interpreted in terms of developmental delay, or precocity. The results here, however, align better with those of Shaw et al. (2006). They reported that more intelligent children show a more prolonged period of cortical thickening, i.e. that peak cortical thickness is obtained later in more intelligent children. Additionally, though IQ is age-adjusted, and tends to be stable across life, individuals who show rapid cortical thinning tend to also show reductions in IQ, particularly PIQ (Burgaleta et al., 2014). In accord with this, the results here are also stronger for PIQ than for VIQ. But, the apparent conflict with the results of Erus et al. (2015) is intriquing. The difference may stem from the differences in the age-range of the subjects, or from the different cognitive measures. The relation between cognitive processing speed and IQ during development is not clear. Or, it may be that residuals based on measures on the cortical surface provide opposing information to those based on measures of deeper structure, including e.g. subcortical gray matter. These are interesting possibilities to investigate in future work.

In sum, we have shown that, contrary to previous results indicating that surface-based measures, e.g. cortical thickness and surface area, relate more weakly to developmental age (Brown et al., 2012; Khundrakpam et al., 2015) than do measures based on subcortical gray or deep white matter, white/gray contrast at the inner edge of cortical gray matter is strongly related to chronological age. Further, our model revealed opposing spatial patterns of predictor importance for contrast and thickness, with primary sensory areas differentiated from association areas, suggesting differential patterns of subcortical connectivity as the basis for the spatial differences, and indicating a strong relation between the two measures. Lastly, we have shown that the residuals of age-prediction based on this white/gray contrast metric are more strongly related to IQ than are those from either thickness or thickness and contrast together, suggesting that this metric is uniquely sensitive to aspects of brain development that reflect cognitive performance.

## 5. Acknowledgments

This research has been supported by the Academy of Finland, and by grant ANRP-MIRI13-3388 from the Azrieli Neurodevelopmental Research Program in partnership with the Brain Canada Multi-Investigator Research Initiative (to ACE). It also benefited from computational resources provided by Compute Canada (www.computecanada.ca) and Calcul Quebec (www.calculquebec.ca).

Data used in preparation of this study were obtained from the Brain Development Cooperative Group and the Pediatric Imaging, Neurocognition, and Genetics Study databases. As such, the investigators within these consortia contributed to the collection of the data, but did not participate in the analysis or writing of this paper. Information related to the Brain Development Cooperative Group can be found at www.nih-pediatricmri.org. Information related to the Pediatric Imaging, Neurocognition, and Genetics Study can be found at http://pingstudy.ucsd.edu.

Thanks to Vladimir Fonov for help with implementation of the contrast measure.

## Footnotes

1 The Matlab code used for constructing stratified cross-validation folds for regression is available at https://github.com/jussitohka/general_matlab

2 The matlab code for both tests is available in www.github.io/jussitohka

## 7. Appendix: Bootstrap confidence intervals for averaged cross-validated correlation coefficient

After 10 CV runs, we have 10 predicted age values *ŷ_i_*(*j*) for each subject *i* (*j* = 1,…, 10). The true age for subject *i* is *y_i_*. For *b* = 1,…, *B*, we

1. sample, with replacement, *N* subjects *I_b_* = [*I_b_*(1),…, *I_b_*(*N*)];
2. for *j* = 1,…, 10, compute *c_bj_* = *corr*(*y_I_b__*,*ŷ_I_b__*(*j*));
3. compute *c_b_* = (1/10) Σ *c_bj_*.

Finally, the 95 % confidence interval is obtained by taking 2.5 % and 97.5 percentiles of {*c_b_* : *b* = 1,…, *B*}.

Above, *corr*(*x*,*y*) denotes the correlation between the vectors *x* and *y*, *y_I_b__* is a shorthand for [*y*_*I*_*b*_(1)_,…, *y*_*I*_*b*_(*N*)_], and *ŷ_I_b__* (*j*) is a shorthand for [*ŷ*_*I*_*b*_(1)_(*j*),…, *ŷ*_*I*_*b*_(*N*)_(*j*)]. The essential point to note is that the same bootstrap samples are applied for every CV run. We note that the permutation test for comparing two MAEs is build upon the same principles and the modification of the above computations for the permutation test is straightforward.

## Supplementary Material

**Figure S1:**
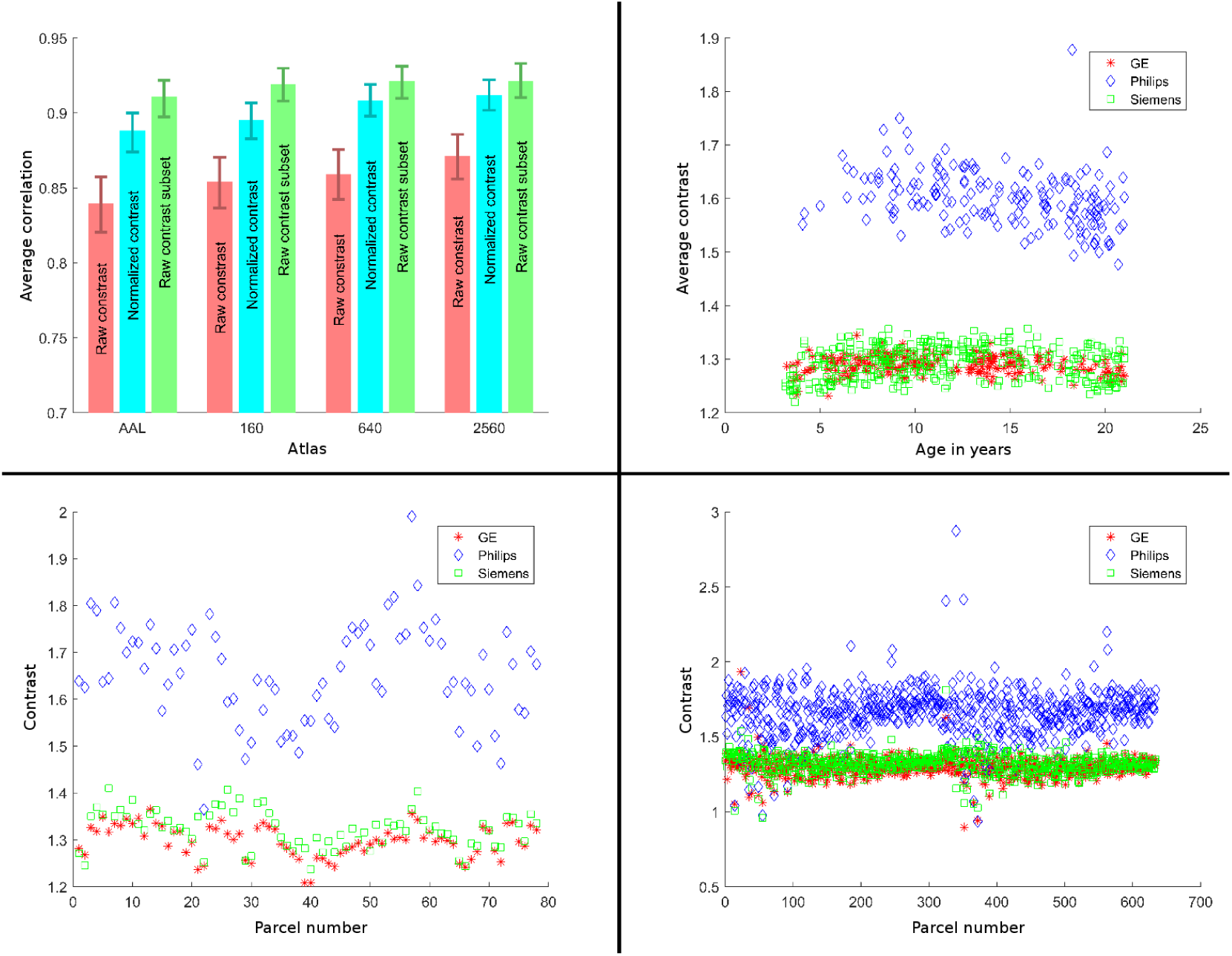
The scanner effect with contrast measures. Upper left panel: Cross-validated average correlations and 95 % CIs with contrast measure a) without scanner manufacturer normalization (red), b) with normalization (cyan) and c) without normalization when considering only the scans acquired using GE or Siemens (i.e. no Philips scans; green). Normalization increased age prediction accuracy and the correlations between predicted and chronological age were almost as high with normalization than with eliminating Philips scans. Upper right panel: Average contrast values across the brain as a function of age. Philips contrast measures were clearly different from other scanners. Lower panels: Examples of contrast profiles across the cortex with three age matched (approximately 16.5 years) girls whose data was acquired using different scanners. Left: AAL parcels and right: 640 parcels. The scanner had a clear effect on the average contrast value and also to its variation across the cortex.

**Figure S2:**
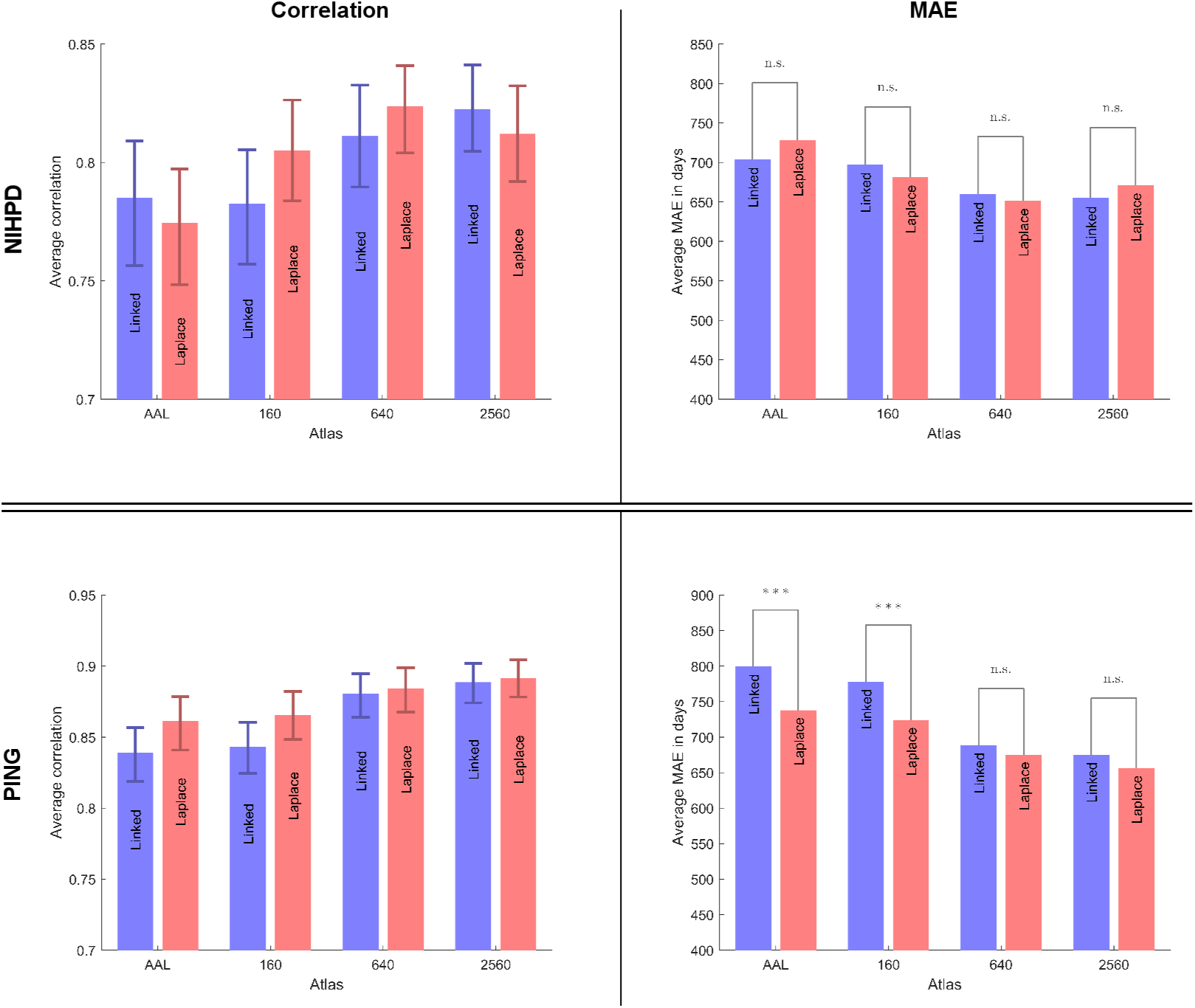
The accuracies of the age prediction using two different cortical thickness measures: linked Euclidean distance and Laplacian distance. Glmnet was used as the machine learning algorithm. See the caption of Fig. 2 for explanation of the plots. The two measures were not significantly different in terms of correlation for either dataset; however, in terms of mean absolute error, while the results for the NIHPD dataset were not significantly different, with the PING dataset, Laplacian distance produced significantly lower errors with some atlases, and never produced higher error. This confirms our choice of the distance measure used in the cortical thickness analysis.

